# EB1 binding provides a diffusion trap mechanism regulating STIM1 localization and Ca^2+^ signaling

**DOI:** 10.1101/224162

**Authors:** Chi-Lun Chang, Yu-Ju Chen, Jen Liou

## Abstract

The endoplasmic reticulum (ER) Ca^2+^ sensor STIM1 forms oligomers and translocates to ER-plasma membrane (PM) junctions to activate store-operated Ca^2+^ entry (SOCE) following ER Ca^2+^ depletion. STIM1 also directly interacts with end binding protein 1 (EB1) at microtubule (MT) plus-ends and resembles comet-like structures during time-lapse imaging. Nevertheless, the role of STIM1-EB1 interaction in regulating SOCE remains unresolved. Using live-cell imaging combined with pharmacological perturbation and a reconstitution approach, we revealed that EB1 binding constitutes a diffusion trap mechanism restricting STIM1 targeting to ER-PM junctions. We further showed that STIM1 oligomers retain EB1 binding ability in ER Ca^2+^-depleted cells. EB1 binding delayed the translocation of STIM1 oligomers to ER-PM junctions and recaptured STIM1 to prevent excess SOCE and ER Ca^2+^ overload. Thus, the counterbalance of EB1 binding and PM targeting of STIM1 shapes the kinetics and amplitude of local SOCE in regions with growing MTs, and contributes to precise spatiotemporal regulation of Ca^2+^ signaling crucial for cellular functions and homeostasis.

**Summary:** STIM1 activates store-operated Ca^2+^ entry (SOCE) by translocating to endoplasmic reticulum-plasma membrane junctions. Chang et al. revealed that STIM1 localization and SOCE are regulated by a diffusion trap mechanism mediated by STIM1 binding to EB1 at growing microtubule ends.

## Introduction

Ca^2+^ is a universal second messenger that governs many important cellular functions, such as secretion, cell migration, differentiation, and apoptosis (Berridge et al., 2000; Dupont et al., 2011; Lewis, 2011). Elevation of cytosolic Ca^2+^ via inositol 1,4,5-triphosphate (IP3)-induced Ca^2+^ release from the endoplasmic reticulum (ER) store following cell surface receptor activation is the key to Ca^2+^ signaling. Animal cells have evolved a feedback mechanism, namely store-operated Ca^2+^ entry (SOCE) that links ER Ca^2+^ store depletion to a Ca^2+^ influx across the plasma membrane (PM) from the extracellular space to support sustained Ca^2+^ signaling and ER Ca^2+^ store refill (Feske and Prakriya, 2013; Prakriya and Lewis, 2015). The importance of SOCE is demonstrated by the patients with mutations in SOCE components manifesting the symptoms of immunodeficiency, autoimmunity, and skeletal myopathy (Feske, 2011).

SOCE is mediated by the ER Ca^2+^ sensor STIM1 and the PM Ca^2+^ channel Orail. The activation of SOCE is a dynamic process involving changes in STIM1 subcellular localization. STIM1 is an ER transmembrane (TM) protein with an N-terminal Ca^2+^-sensing EF hand-SAM (EF-SAM) domain in the ER lumen (Figure 1A). The cytosolic portion of STIM1 contains coiled-coil domains (CC1 to CC3), a serine/proline (S/P) region, and a C-terminal region (CT, amino acid 633-685, Figure 1A) with a polybasic motif (PB). In the resting state, STIM1 binds to Ca^2+^ in the ER lumen and localizes diffusely throughout the ER (Liou et al., 2005). Following ER Ca^2+^ store depletion, Ca^2+^-free STIM1 rapidly oligomerizes leading to a conformational extension of the PB and the Orai1 activation domain, namely CAD, SOAR, or CC9 that roughly corresponds to the CC2-CC3 domains (Kawasaki et al., 2009; Park et al., 2009; Prakriya and Lewis, 2015; Yuan et al., 2009). The oligomerized/exposed PB binds to phosphatidylinositol 4,5-bisphosphate (PIP2) at the PM (Ercan et al., 2009; Liou et al., 2007; Walsh et al., 2010). STIM1-PIP2 interaction traps STIM1 at ER-PM junctions where the ER and the PM form close appositions allowing STIM1 at the ER to activate Orai1 at the PM resulting in SOCE. STIM1 targeting to ER-PM junctions is a rate-limiting step in the activation of SOCE. Although STIM1 oligomerization occurs within 5 s following ER Ca^2+^ store depletion, it takes more than 40 s for STIM1 to translocate to ER-PM junctions (Liou et al., 2007). The mechanism underlying the time discrepancy between STIM1 oligomerization and translocation is not clear.

**Figure 1.**
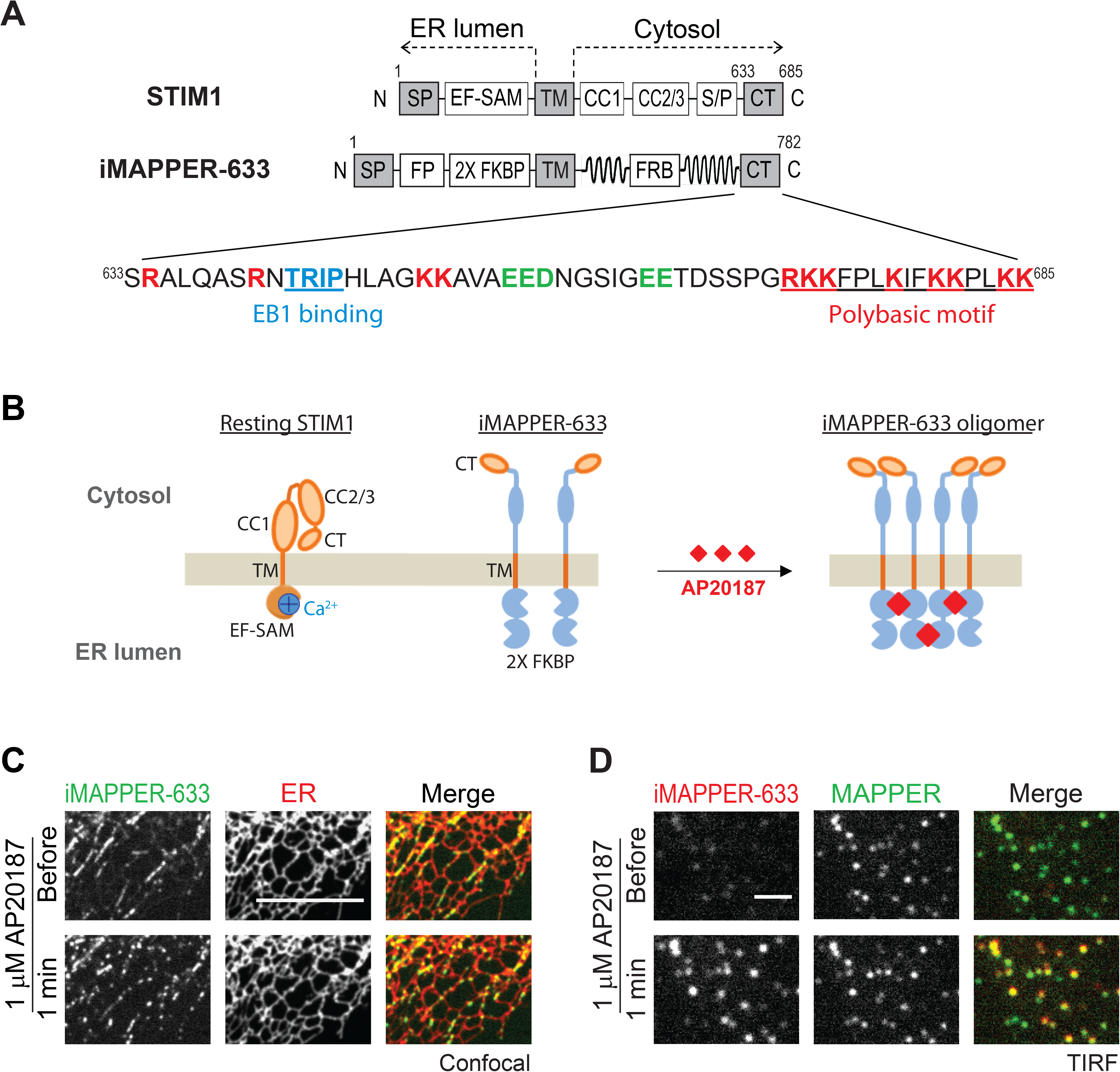
iMAPPER-633 Recapitulates Dynamic STIM1 Subcellular Localization. (A) Diagrams of STIM1 and iMAPPER-633. Amino acid number and domains are indicated. SP, signal peptide; EF-SAM, EF hand and sterile alpha motif; TM, transmembrane domain; CC1-CC3, coiled coil domain 1-3; S/P, serine and proline rich region; CT, C-terminal (633-685); FP, fluorescence protein; 2X FKBP, tandem FKBP domain; FRB, FKBP-rapamycin binding domain. Identical domains between STIM1 and iMAPPER-633 are in gray. The amino acid sequences of STIM1 CT are displayed. Core EB1 binding motif are labeled in blue. Positively charged residues are labeled in red. Negative charged residues are labeled in green. (B) A schematic diagram depicting STIM1 and iMAPPER-633 at basal and oligomerized iMAPPER-633 following AP20187 treatment. Domains are indicated as in (A). (C) YFP-iMAPPER-633 displays punctate localization following 1 μM AP20187 treatment monitored by confocal microscopy in HeLa cells co-transfected with mCherry-ER. Scale bar, 10 μm. (D) Translocation of mCherry-iMAPPER-633 to ER-PM junctions following 1 μM AP20187 treatment monitored by TIRF microscopy in HeLa cells co-transfected with GFP-MAPPER. Scale bar, 2 μm.

In addition to PIP2 and Orail binding, STIM1 directly interacts with the microtubule (MT)-plus-end binding proteins EB1 and EB3 to a lesser extent (Grigoriev et al., 2008). These interactions are mediated by an EB1 binding motif that resides in the CT. STIM1 interaction with EB1 at MT plus-ends can be visualized using live-cell imaging as overexpressed STIM1 adopts MT-like organization and displays comet-like structures (Baba et al., 2006; Grigoriev et al., 2008; Honnappa et al., 2009; Mercer et al., 2006). STIM1 comets represent sub-populations of STIM1 that are transiently trapped by binding to EB1 at the contacts between MT plus-ends and the ER (Grigoriev et al., 2008). Mutation of the core EB1 binding TRIP residues to TRNN disrupts STIM1-EB1 interaction, resulting in the disappearance of STIM1 comets (Honnappa et al., 2009). Nevertheless, the significance of STIM1 trapping by EB1 binding in regulating STIM1 translocation to ER-PM junctions and SOCE remains unclear.

To dissect the contribution of EB1 binding to STIM1 localization and function, we generated iMAPPER-633, a mini-STIM1 construct that contains the ER targeting motifs and the CT (amino acid 633-685), and lacks the EF-SAM, coiled-coil domains and the S/P region to mediate Ca^2+^ sensing, Orai1 binding and phosphorylation, respectively. iMAPPER-633 binds to EB1 at MT plus-ends and translocates to ER-PM junctions following chemically-induced oligomerization, indicating that it contains the necessary components to recapitulate STIM1 localization in the resting state and during ER Ca^2+^ store depletion. Experiments using iMAPPER-633 also revealed that EB1 binding is dominant over PIP2 binding within the CT and that oligomerization strongly potentiates PIP2 binding, resulting in translocation. We further showed that EB1 binding restricts STIM1 diffusion in the ER and limits STIM1 access to ER-PM junctions in the resting state and during ER Ca^2+^ store depletion. Disruption of EB1 binding facilitates Orai1 recruitment and SOCE activation, resulting in ER Ca^2+^ overload. Together, our findings indicate that EB1 binding provides a diffusion trap mechanism regulating STIM1 localization and SOCE, and suggest that STIM1-mediated Ca^2+^ signaling may be locally regulated by binding to EB1 on growing MT ends.

## Results

### iMAPPER-633: a mini-STIM1 construct that recapitulates dynamic STIM1 subcellular localization

The localization of STIM1 is regulated by multiple factors including ER Ca^2+^ levels, oligomerization, phosphorylation, as well as binding to Orai1, PIP2, and EB1. The role of EB1 binding in regulating STIM1-mediated Ca^2+^ signaling at ER-PM junctions is not well understood. To dissect the contribution of EB1 binding in STIM1 translocation to ER-PM junctions, we employed a reconstitution approach and engineered a mini-STIM1 construct that contains the signal peptide (SP) and TM of STIM1 for ER membrane localization, as well as the CT of STIM1 enabling its binding to EB1 and PIP2 (Figure 1A). A tandem FKBP (FK506 binding protein) motif (2X FKBP) was inserted into the ER luminal region following a fluorescence protein (FP) to enable oligomerization upon treatment of a small molecule AP20187, and optical imaging, respectively. (Figure 1B). We further added in the cytosolic linker region of the synthetic ER-PM junctional marker MAPPER (Chang et al., 2013), which has been shown to provide the proper length spanning the gap at ER-PM junctions. We named this mini-STIM1 construct “iMAPPER-633” (inducible MAPPER-633) because its design resembles MAPPER and features inducible translocation to ER-PM junctions via the CT of STIM1 starting at residue 633. An intermediate construct containing the SP, an FP, the 2X FKBP, and the TM displayed ER localization when expressed in HeLa cells, indicating successful ER targeting (Figure S1A). Unlike the CT in full-length STIM1 which is partially buried in the resting state (Zhou et al., 2013), the CT in iMAPPER-633 is expected to be fully exposed, facilitating assessment of the contribution of EB1 -binding and PIP2-binding motifs to STIM1 localization (Figure 1B).

When iMAPPER-633-transfected HeLa cells were examined under confocal microscopy, iMAPPER-633 displayed comet-like structures moving toward the cell periphery, similar to resting STIM1 that binds to EB1 at MT plus-ends (upper panels in Figure 1C and Movie S1). These iMAPPER-633 comets colocalized with an ER luminal marker and with STIM1 (Figures 1C and S1B), suggesting that iMAPPER-633 is an ER protein concentrated at the contacts between MT plus-ends and the ER. Consistent with the expectation that the CT of iMAPPER-633 is fully exposed and more accessible to EB1 binding than that of STIM1, iMAPPER-633 appeared to be more concentrated in MT-like structures than in the ER compared to STIM1 (Figure S1B). The extensive trapping of iMAPPER-633 at MT-like structures was accompanied by a few highly concentrated iMAPPER-633 clusters with apparent movement toward the nucleus, possibly formed due to loss of EB1 binding during MT catastrophe (Figure S1B yellow arrowheads). When AP20187 was applied to induce oligomerization, iMAPPER-633 rapidly translocated into puncta while the bulk ER structure was unaffected (lower panels in Figure 1C). Colocalization of iMAPPER-633 with MAPPER, monitored by total internal reflection fluorescence (TIRF) microscopy, indicates that iMAPPER-633 puncta formation corresponds to its translocation to ER-PM junctions (Figure 1D). These results further suggest that iMAPPER-633 contains targeting motifs sufficient for recapitulating the subcellular localization of STIM1 in the resting state as well as in the oligomized state induced by ER Ca^2+^ store depletion. These results also revealed that the exposed CT of STIM1 preferentially binds to EB1 at MT plus-ends rather than PIP2 at the PM, and that oligomerization strongly potentiates PIP2 binding, resulting in translocation.

### Inhibition of EB1 binding triggers iMAPPER-633 translocation to ER-PM junctions

Next, we applied nocadazole (NocZ), an inhibitor of MT polymerization, to disrupt EB1-MT association. We found that iMAPPER-633 translocated to ER-PM junctions following NocZ treatment as EB1 comets disappeared (Figure 2A). Consistently, we observed that iMAPPER-633 readily localized at ER-PM junctions in cells with EB1 knockdown by small interfering RNA against EB1 (siEB1) (Figures 2B and 2C). We further generated the iMAPPER-633-TRNN mutant incapable of EB1 binding and found that iMAPPER-633-TRNN pre-localized to ER-PM junctions (upper panels in Figure 2D). Notably, the intensity of iMAPPER-633-TRNN puncta remained similar following AP20187 treatment, suggesting that the majority of iMAPPER-633-TRNN was trapped at ER-PM junctions as a result of its inability of EB 1 binding (lower panels in Figure 2D). These data indicate that the PB in the exposed CT is sufficient for trapping iMAPPER-633 at ER-PM junctions and that EB1 binding diverts iMAPPER-633 to be trapped at MT plus-ends, preventing its localization at ER-PM junctions.

**Figure 2.**
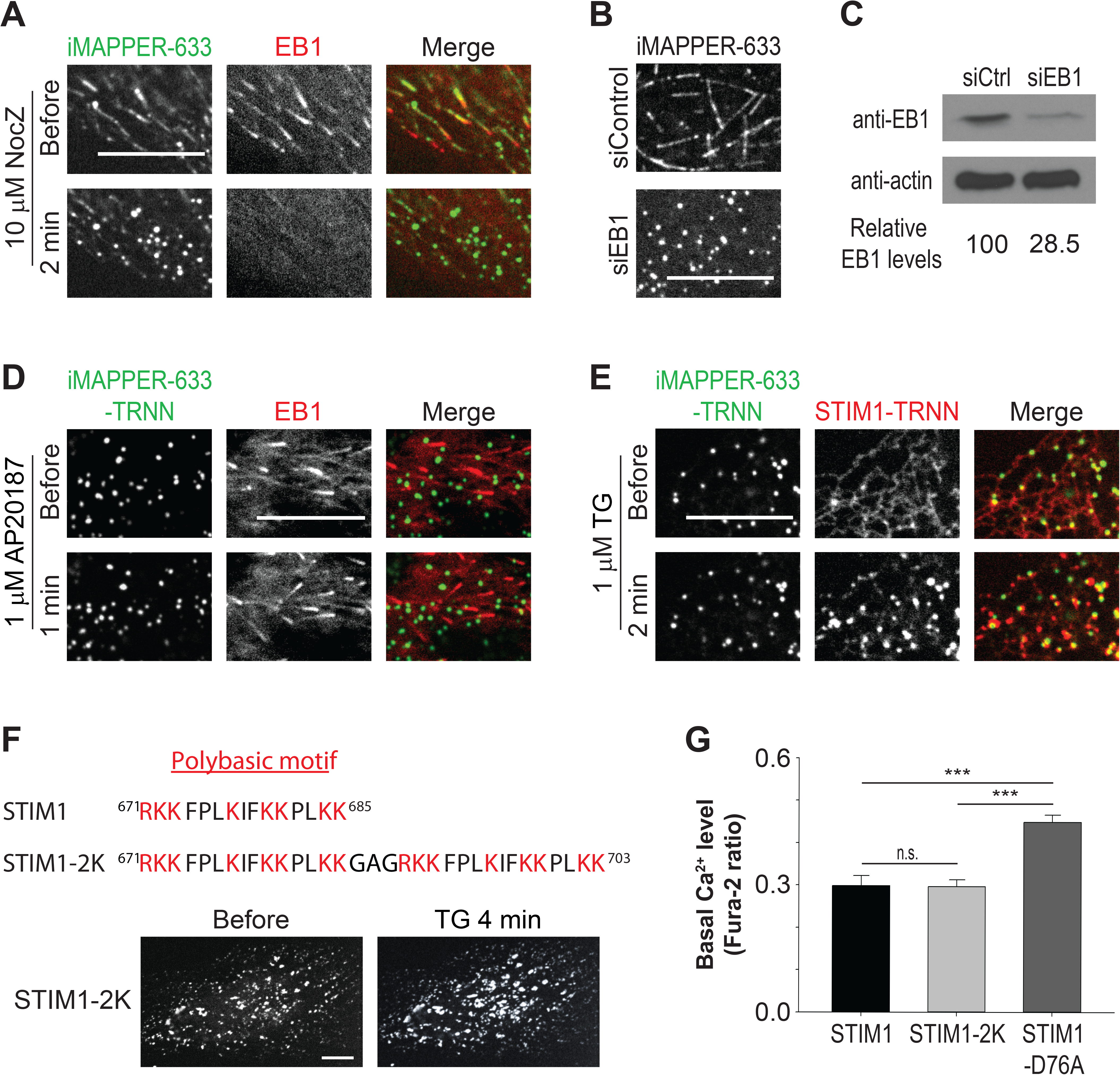
Inhibition of EB1 Binding Triggers iMAPPER-633 Translocation to ER-PM Junctions. (A) Translocation of YFP-iMAPPER-633 to ER-PM junctions following 10 μM NocZ treatment monitored by confocal microscopy in HeLa cells co-transfected with EB1-mCherry. Scale bar, 10 μm. (B) Subcellular localizations of YFP-iMAPPER-633 monitored by confocal microscopy in HeLa cells transfected with siControl or siEB1. Scale bar, 10 μm. (C) EB1 protein levels detected by western blotting using anti-EB1 antibody in HeLa cells transfected with siControl (siCtrl) or siEB1. The intensity of bands was measured by image J. Relative EB1 levels are indicated. (D) YFP-iMAPPER-633-TRNN distributes to ER-PM junctions in the absence or presence of AP20187 in HeLa cells monitored by confocal microscopy. Scale bar, 10 μm. (E) Translocation of mCherry-STIM1-TRNN to ER-PM junctions labeled by YFP-iMAPPER-633-TRNN following 1 μM TG treatment in HeLa cells monitored by confocal microscopy. Scale bar, 10 μm. (F) mCherry-STIM1-2K with two PB in tandem in the CT distributes to ER-PM junctions in the absence or presence of 1 μM TG in HeLa cells monitored by confocal microscopy. Scale bar, 10 μm. (G) Basal cytosolic Ca^2+^ levels monitored by Fura-2 ratio in HeLa cells transfected with mCherry-STIM1, mCherry-STIM1-2K, or mCherry-STIM1-D76A. Mean ± SD are shown (3 independent experiments). n.s., not significant; ***, *p* < 0.001.

Unlike iMAPPER-633-TRNN, the STIM1-TRNN mutant showed a uniform distribution throughout the ER with minimal pre-localization at ER-PM junctions and no comet-like structures (Figure 2E). ER Ca^2+^ store depletion by thapsigargin (TG) was required to trigger STIM1-TRNN translocation to ER-PM junctions to co-localize with iMAPPER-633-TRNN. These results are consistent with a previous finding that the CT in full-length STIM1 is partially buried and is incapable of binding to PIP2 in the PM until STIM1 activation following ER Ca^2+^ depletion (Zhou et al., 2013). We further generated a STIM1-2K construct by adding an extra PB to the very C-terminus of STIM1 to enhance its PM binding affinity. This STIM1-2K construct with two PB in tandem in the CT pre-localized to ER-PM junctions without ER Ca^2+^ store depletion (Figure 2F). Similar pre-localization has been observed with the STIM1-D76A mutant that contains a point mutation disrupting its ability to bind ER Ca^2+^ and exhibits an active conformation (Liou et al., 2005). Unlike STIM1-D76A, expression of STIM1-2K did not increase basal Ca^2+^ levels (Figure 2G), suggesting that STIM1-2K is not in an active conformation. These results indicate that increased PM binding affinity can cause STIM1 trapping at ER-PM junctions without activating SOCE.

### EB1 binding constitutes a diffusion trap limiting STIM1 localization at ER-PM junctions

By binding to EB1 at MT plus-ends, STIM1 is transiently trapped at ER regions in contact with growing MT end. To examine the effect of EB1 binding on STIM1 diffusion, we performed fluorescence recovery after photobleaching (FRAP) experiments using cells transfected with STIM1 or STIM1-TRNN. We found that STIM1-TRNN fluorescence recovered faster than that of STIM1 in the bleached regions with a significant difference in the time required to reach the half recovery (*t_1/2_*) (Figures 3A and 3B). These results demonstrate that STIM1-EB1 interaction restricts STIM1 diffusion in the ER membrane. Trapping by EB1 at MT plus-ends likely limits the amount of STIM1 molecules accessing other ER regions including ER-PM junctions. To test this hypothesis, we monitored ER-PM junctions in cells co-transfected with a mCherry-tagged ER luminal marker and YFP-tagged STIM1 using TIRF microscopy. Following NocZ treatment, an increase in the intensity ratio of STIM1 over the ER marker at ER-PM junctions was observed, whereas the intensity ratio of STIM1-TRNN over the ER marker at ER-PM junctions remained the same (Figures 3C, 3D and 3E). These findings indicate that NocZ treatment released the sub-population of STIM1 trapped by EB1 at MT plus-ends, resulting in an increase of STIM1 molecules diffusing through ER-PM junctions in resting cells.

**Figure 3.**
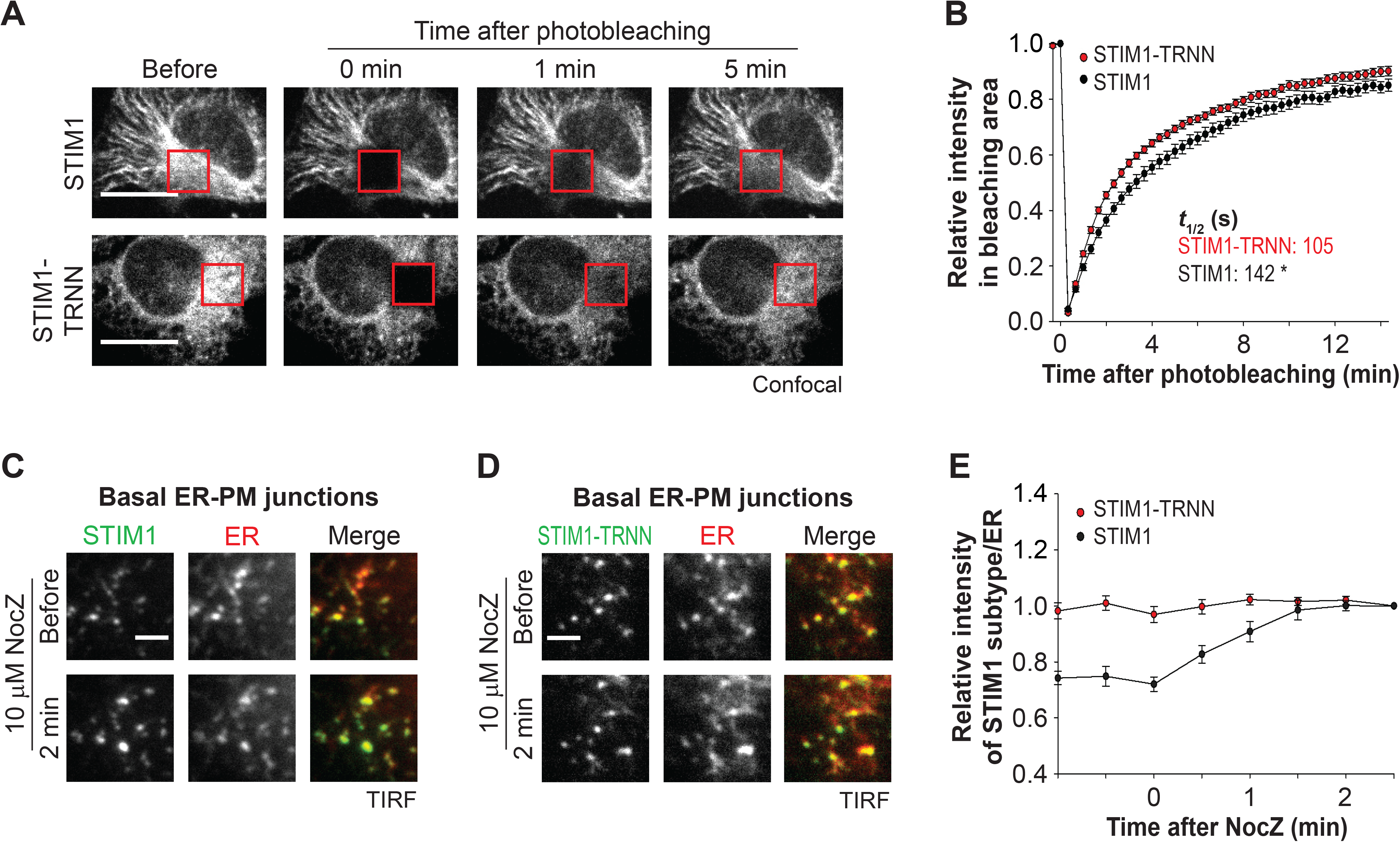
EB1 Binding Constitutes a Diffusion Trap Limiting STIM1 Localization at ER-PM Junctions. (A) Fluorescence recovery of YFP-STIM1 and YFP-STIM1-TRNN after photobleaching (red square boxes) in HeLa cells monitored by confocal microscopy. Scale bar, 10 μm.(B) Relative intensity of YFP-STIM1 and YFP-STIM1-TRNN in the bleached areas as described in (A). Mean ± SEM are shown (19 to 20 cells from 3 independent experiments). Mean times to the half recovery (*t_1/2_*) are indicated. *, *p* < 0.05. (C and D) Changes in intensity of YFP-STIM1 (C) and YFP-STIM1-TRNN (D) at ER-PM junctions following 10 μM NocZ treatment monitored by TIRF microscopy in HeLa cells co-transfected with mCherry-ER. Scale bar, 2 μm. (E) Relative changes in intensity of STIM1 subtypes normalized to the intensity of mCherry-ER as described in (C and D). Mean ± SEM are shown (13 to 14 cells from 3 to 4 independent experiments).

### Activated STIM1 retains EB1 binding ability in ER Ca^2+^-depleted cells

A previous report suggested that STIM1 oligomerization following ER Ca^2+^ store depletion does not preclude its association with endogenous EB proteins (Grigoriev et al., 2008). In support of this idea, we observed partial co-localization of STIM1 and EB1 after TG treatment (Figure 4A). Consistently, immuneprecipitation (IP) experiments showed that a portion of mCherry-STIM1 remained bound to EB1-GFP following TG treatment while mCherry-STIM1-TRNN showed negligible interaction with EB1-GFP (Figures 4B and S2A). We further applied 100 μM ML-9, which has been shown to rapidly trigger STIM1 dissociation from ER-PM junctions and reversion of puncta formation (Smyth et al., 2008), to cells co-transfected with STIM1 and EB1 during ER Ca^2+^ store depletion. Following ML-9 treatment, TG-induced STIM1 puncta rapidly disappeared and STIM1-trapping by EB1 became apparent without refilling the ER Ca^2+^ store (Figure 4C). The disappearance of STIM1 puncta induced by ML-9 treatment was not due to the disruption of ER-PM junctions as monitored by an ER luminal marker using TIRF microscopy (Figure 4D). It is possible that ML-9 abolishes STIM1 trapping by PM PIP2 since STIM1 trapping at ER-PM junctions by Orai1 was not affected by ML-9 treatment (Figure S2B). Furthermore, we found that STIM1-D76A, a constitutively active mutant that pre-localizes at ER-PM junctions without ER Ca^2+^ store depletion, was immediately trapped by EB1 following ML-9 treatment (Figure 4E; Movie S2), whereas STIM1-D76A-TRNN displayed ER localization after ML-9-induced dissociation from ER-PM junctions (Figure 4F). Together, these results demonstrate that STIM1 can bind to EB1 at MT plus-ends regardless of its activation state and the level of ER Ca^2+^ store. These findings indicate that EB1 at MT plus-ends can still capture and trap STIM1 during ER Ca^2+^ depletion.

**Figure 4.**
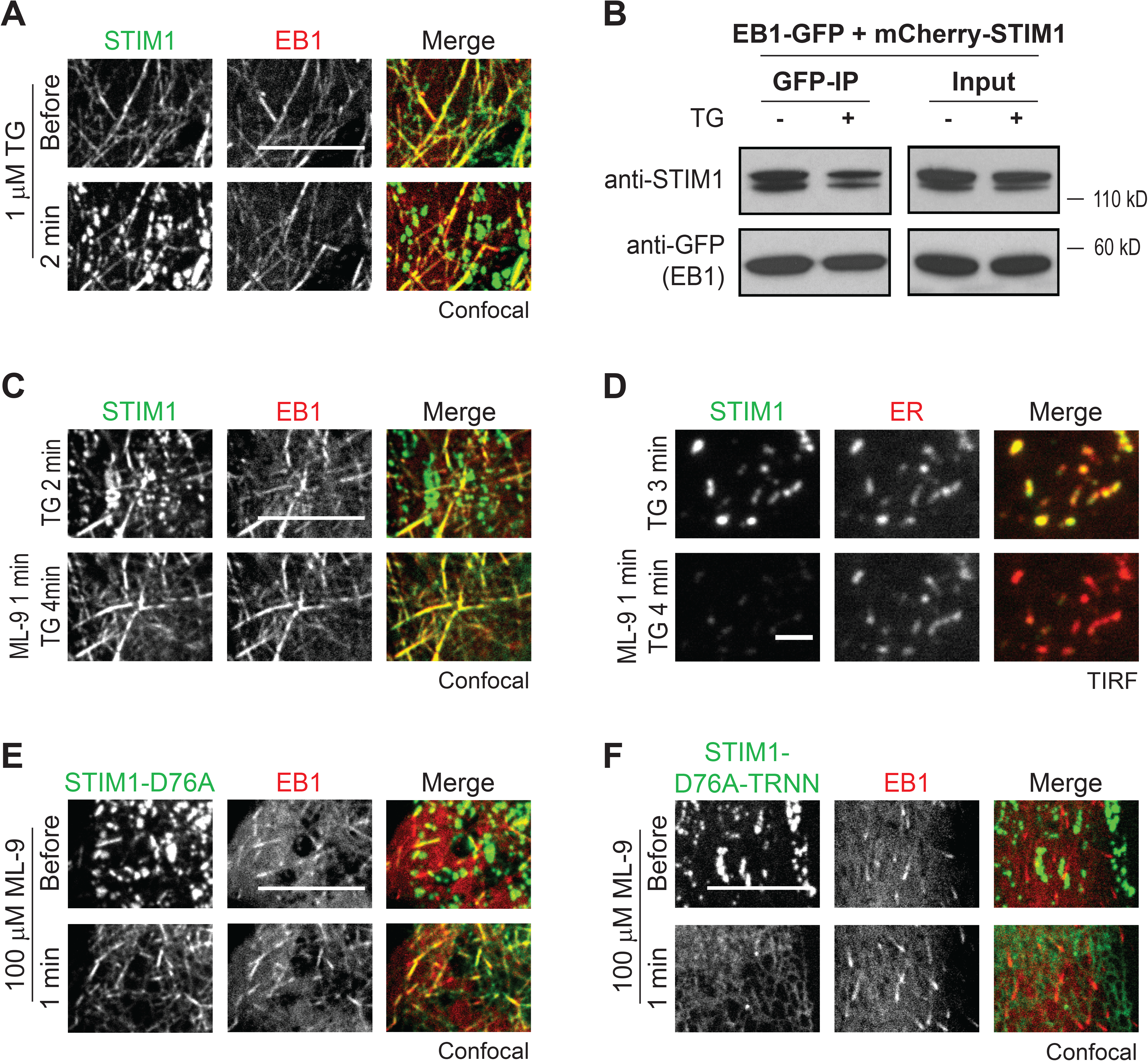
Activated STIM1 Retains EB1 Binding Ability in ER Ca^2+^-depleted Cells. (A) Localization of YFP-STIM1 and EB1-mCherry in a HeLa cells during the resting state and following 1 μM TG treatment monitored by confocal microscopy. Scale bar, 10 μm.(B) Immunoprecipitation (IP) of EB1-GFP with mCherry-STIM1 following 1 μM TG treatment in HeLa cells. Protein levels of EB1-GFP and mCherry-STIM1 in total cell lysates (Input) and in IP were assessed by western blotting using antibodies against GFP (anti-GFP) and STIM1 (anti-STIM1). (C) Colocalization of YFP-STIM1 and EB1-mCherry in HeLa cells following 100 μM ML-9 treatment during ER Ca^2+^ depletion by 1 μM TG monitored by confocal microscopy. Scale bar, 10 μm. (D) TG-induced YFP-STIM1 puncta disappear at ER-PM junctions labeled by mCherry-ER in HeLa cells following 100 μM ML-9 treatment monitored by TIRF microscopy. Scale bar, 2 μm. (E) Colocalization of YFP-STIM1-D76A and EB1-mCherry in HeLa cells following 100 μM ML-9 treatment monitored by confocal microscopy. Scale bar, 10 μm. (F) YFP-STIM1-D76A-TRNN display the ER localization without colocalizing with EB1-mCherry in HeLa cells following 100 μM ML-9 treatment monitored by confocal microscopy. Scale bar, 10 μm.

### EB1 binding impedes STIM1 translocation to ER-PM junctions and Orai1 recruitment during ER Ca^2+^ depletion

We then reasoned that the diffusion trap mechanism mediated by EB1 binding may impede STIM1 translocation to ER-PM junctions following ER Ca^2+^ store depletion. Consistent with this notion, we observed nearly complete translocation of YFP-STIM1-TRNN 30 s after 1 μM ionomycin treatment while YFP-STIM1 only began to accumulate at ER-PM junctions (Figure 5A). The t1/2 of STIM1 translocation to ER-PM junctions was 56.2 s (Figure 5B), which is comparable to a previous report (Liou et al., 2007). By contrast, STIM1-TRNN showed a significantly faster translocation than STIM1 with a t1/2 of 22.5 s. A higher amplitude of STIM1-TRNN translocation to ER-PM junctions than that of STIM1 was observed, indicating enhanced accumulation of STIM1-TRNN compared to STIM1 at ER-PM junctions (Figure 5B). The kinetic differences in translocation to ER-PM junctions between STIM1 and STIM1-TRNN were also detected in cells treated with TG (Figures S3A and S3B). Consistently, a significant increase in the rate of STIM1 translocation following ionomycin treatment was detected in siEB1-treated cells compared with those treated with siControl (Figures S3C and S3D). Notably, disruption of STIM1-EB1 interaction consistently led to an accelerated STIM1 translocation by 23-34 s regardless of the rate of ER Ca^2+^ store depletion. These results indicate that trapping by EB1 at MT plus-ends delays diffusion of STIM1 oligomers to ER-PM junctions during ER Ca^2+^ store depletion.

**Figure 5.**
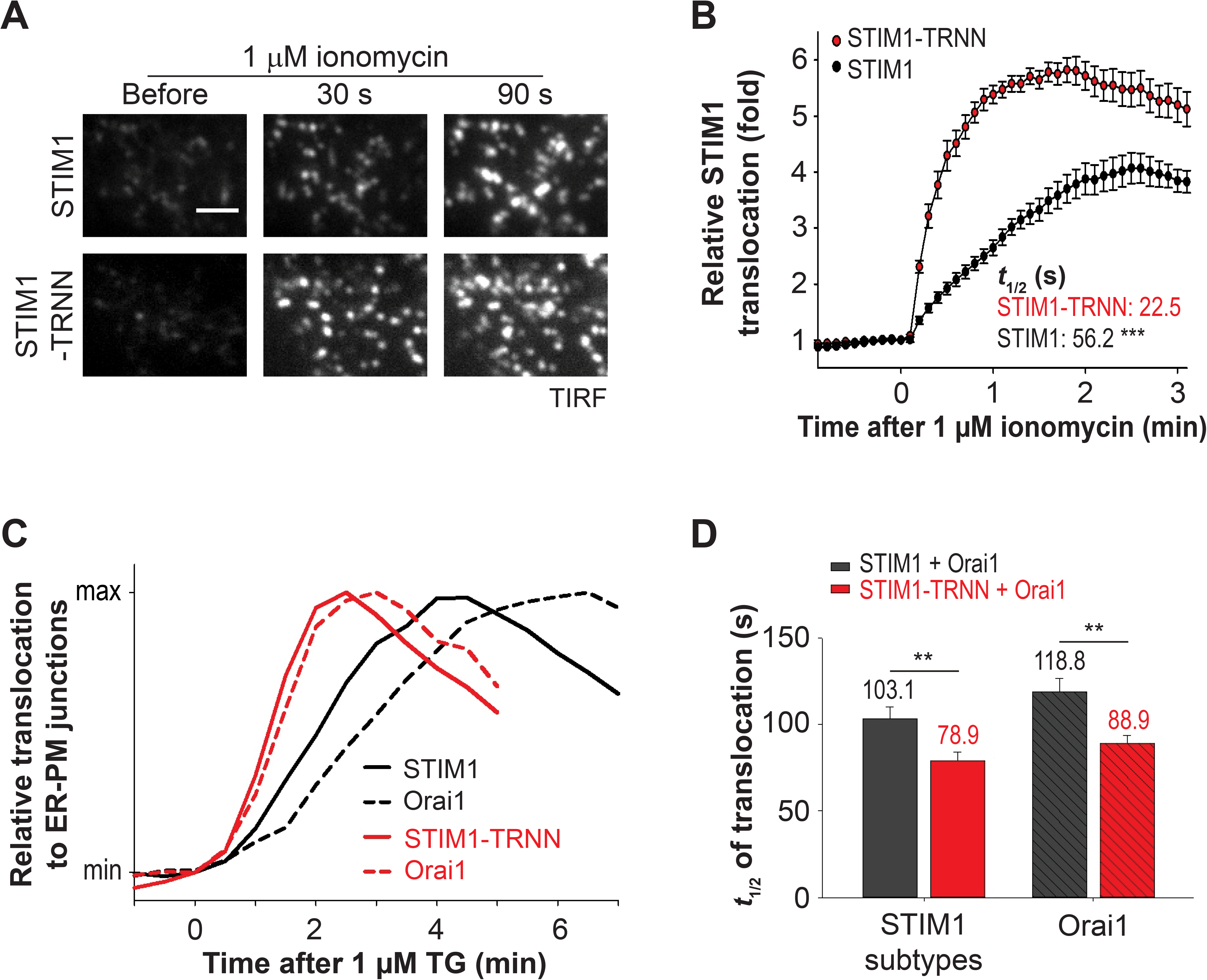
EB1 Binding Impedes STIM1 Translocation to ER-PM Junctions and Orai1 Recruitment during ER Ca^2+^ Depletion. (A) Translocation of YFP-STIM1 and YFP-STIM1-TRNN to ER-PM junctions following 1 μM ionomycin treatment in HeLa cells monitored by TIRF microscopy. Scale bar, 2 μm. (B) Relative translocation to ER-PM junctions of YFP-STIM1 and YFP-STIM1-TRNN as described in (A). Mean ± SEM are shown (14 to 15 cells from 3 independent experiments). Mean times to the half-maximal translocation (*t_1/2_*) are indicated. ***, *p* < 0.001. (C) Relative translocation to ER-PM junctions of YFP-STIM1 subtypes and corresponding Orail-mCherry following 1 μM TG treatment in HeLa cells monitored by TIRF microscopy. Black: YFP-STIM1 co-expressed with Orai1-mCherry; Red: STIM1-TRNN co-expressed with Orai1-mCherry. Mean traces are shown (15 to 23 cells from 3 to 4 independent experiments). (D) Times to the half-maximal translocation of YFP-STIM1 subtypes and Orai1-mCherry as described in (C). Mean ± SEM are shown. **, *p* < 0.01.

Activated STIM1 trapped at ER-PM junctions can bind to the PM Ca^2+^ channel Orai1, resulting in Orai1 recruitment to ER-PM junctions. We further monitored Orai1-mCherry translocation to ER-PM junctions in cells co-expressing YFP-STIM1 or YFP-STIM1-TRNN. Following TG treatment, translocation of STIM1 and STIM1-TRNN preceded Orai1 recruitment to ER-PM junctions with a difference in t1/2 of 15 s (103.1 s for STIM1 and 118.8 s for Orai1) and 10 s (78.9 s for STIM1-TRNN and 88.9 s for Orai1), respectively (Figures 5C and 5D). Notably, a significant acceleration of Orai1 accumulation at ER-PM junctions by 30 s (88.9 s vs. 118.8 s) was observed in STIM1-TRNN overexpressing cells compared with STIM1-overexpressing one (Figure 5D). These data indicate that STIM1-trapping by EB1 at MT plus-ends delays the binding and recruitment of Orai1 by STIM1 at ER-PM junctions during ER Ca^2+^ store depletion.

### Disruption of EB1 binding facilitated SOCE and resulted in ER Ca^2+^ store overload

STIM1 interaction with Orai1 at ER-PM junctions initiates SOCE. Thus, regulation of STIM1 localization by EB1 binding in the resting state and during ER Ca^2+^ store depletion likely shapes the dynamics and extent of SOCE. Consistent with this notion, the sustained phase of cytosolic Ca^2+^ levels following ER Ca^2+^ depletion induced by histamine and TG treatment in siEB1-treated cells was higher than that in siControl-transfected cells, suggesting an enhanced SOCE (Figure 6A). Next, we selectively monitored Ca^2+^ entry from the extracellular space following ER Ca^2+^ store depletion as a specific readout for SOCE and found that knockdown of EB1 resulted in a significant increase in the amplitude of SOCE (Figure 6B). Consistently, STIM1-TRNN overexpression also led to a significant enhancement of SOCE than that mediated by STIM1 overexpression (Figure 6C). The effect of EB1-mediated STIM1 diffusion trap on Ca^2+^ signaling was further exemplified in experiments using the constitutively active STIM1-D76A and STIM1-D76A-TRNN constructs. Overexpression of STIM1-D76A led to a marked increase in basal cytosolic Ca^2+^ levels compared to control (STIM1-D76A vs. TM, Figure 6D). This increase is due to a constitutive SOCE because removal of extracellular Ca^2+^ rapidly decreased the cytosolic Ca^2+^ to the control level and re-addition of extracellular Ca^2+^ rapidly restored the elevated cytosolic Ca^2+^ level in STIM1-D76A overexpressing cells. Overexpression of STIM1-D76A-TRNN further potentiates the elevated cytosolic Ca^2+^ level at basal and following re-addition of extracellular Ca^2+^ (Figure 6D). Together, these results indicate that EB1 binding limits STIM1 localization at ER-PM junctions, dampening the amplitudes of STIM1-mediated SOCE.

**Figure 6.**
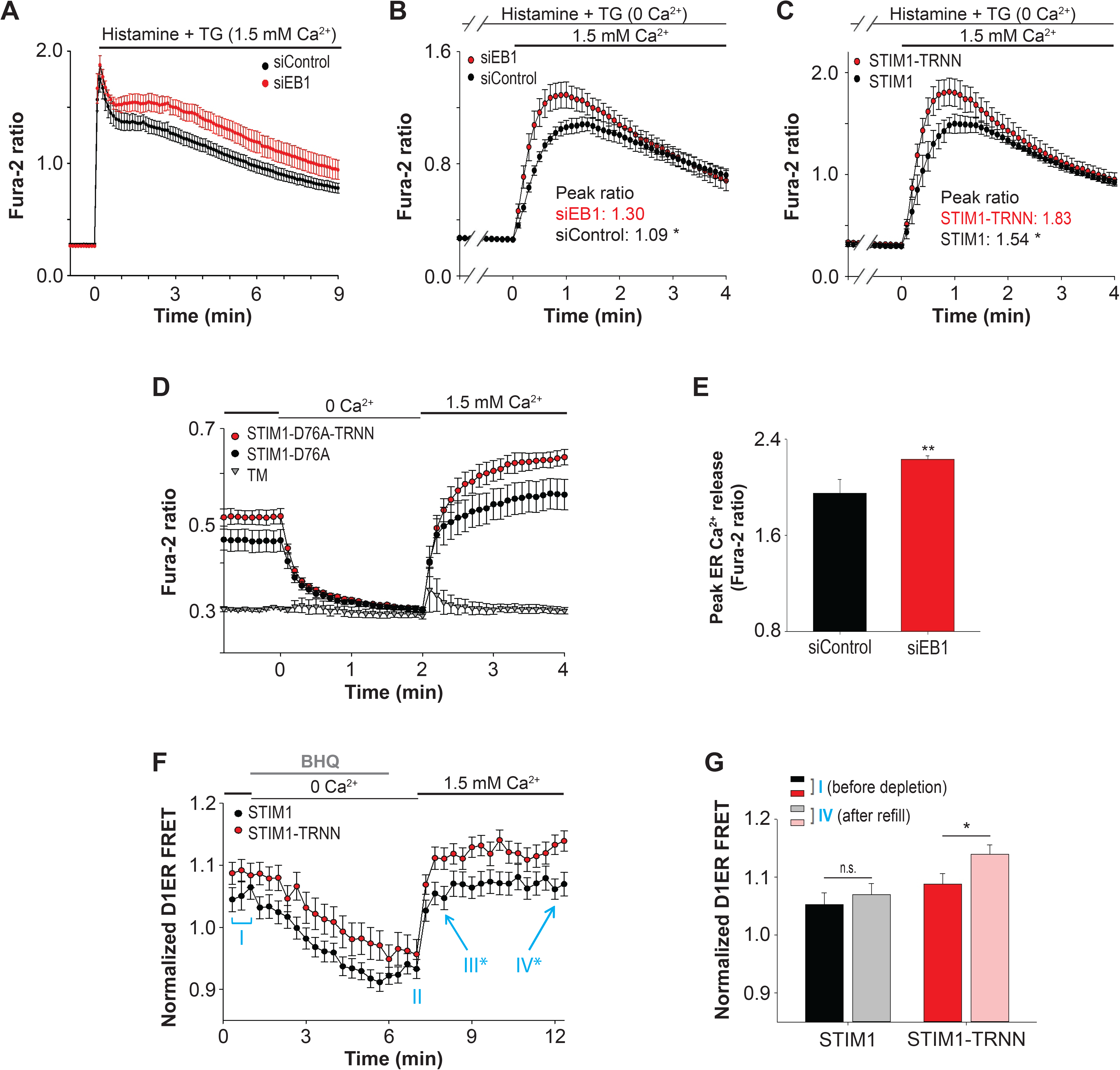
Disruption of EB1 Binding Facilitated SOCE and Resulted in ER Ca^2+^ Store Overload. (A) Relative changes in cytosolic Ca^2+^ concentration following 100 μM histamine and 1 μM TG treatment monitored by Fura-2 ratio in HeLa cells transfected with siControl or siEB 1. Mean ± SEM are shown (4 independent experiments). (B) SOCE triggered by 100 μM histamine and 1 μM TG treatment monitored by Fura-2 ratio in HeLa cells transfected with siControl or siEB 1. Mean ± SEM are shown (3 independent experiments). Peaks of Fura-2 ratio are indicated. *, *p* < 0.05. (C) SOCE triggered by 100 μM histamine and 1 μM TG treatment monitored by Fura-2 ratio in HeLa cells transfected with YFP-STIM1 or YFP-STIM1-TRNN. Mean ± SEM are shown (3 independent experiments). Peaks of Fura-2 ratio are indicated. *, *p* < 0.05. (D) Relative changes in cytosolic Ca^2+^ concentration in response to depletion and re-addition of extracellular Ca^2+^ monitored by Fura-2 ratio in HeLa cells transfected with YFP-TM, YFP-STIM1-D76A, or YFP-STIM1-D76A-TRNN. Mean ± SEM are shown (3 to 4 independent experiments). (E) Peak ER Ca^2+^ release by 1 μM ionomycin treatment in the absence of extracellular Ca^2+^ monitored by Fura-2 ratio in HeLa cells transfected with siControl or siEB 1. Mean ± SEM are shown (3 independent experiments). **, *p* < 0.01. (F) Relative ER Ca^2+^ levels in the resting state (phase I), following 5 μM BHQ treatment (phase II), and after BHQ washout (phase III and IV) monitored by D1ER in HeLa cells transfected with mCherry-STIM1 or mCherry-STIM1-TRNN. Mean ± SEM are shown (16 to 26 cells from 3 independent experiments). *, *p* < 0.05 between STIM1 and STIM1-TRNN. (G) Relative ER Ca^2+^ levels in phase I and IV as described in (F) monitored by D1ER in HeLa cells transfected with mCherry-STIM1 or mCherry-STIM1-TRNN. Mean ± SEM are shown. n.s., not significant; *, *p* < 0.05.

A main function of SOCE is to refill the ER Ca^2+^ store following depletion. We further tested if enhanced SOCE caused by the absence of STIM1-EB1 interaction results in ER Ca^2+^ store overload. In the resting cells, we observed a significantly elevated ER Ca^2+^ store in siEB1-treated cells as monitored by the release of ER Ca^2+^ by ionomycin treatment in the absence of extracellular Ca^2+^ (Figure 6E). We further tracked the dynamic changes in ER Ca^2+^ levels during store depletion and refill using an ER Ca^2+^ sensor D1ER (Palmer et al., 2004) and a reversible SERCA inhibitor named BHQ. We found that the level of ER Ca^2+^ was moderately elevated in STIM1-TRNN-transfected cells compared with that in STIM1-transfected cells before and after depletion by BHQ (Figure 6F, phases I and II). Following BHQ washout and re-addition of extracellular Ca^2+^, a significant elevation in the ER Ca^2+^ level was observed in STIM1-TRNN-overexpressing cells compared with that in STIM1-overexpressing cells (Figure 6F, phase III and IV). Notably, the ER Ca^2+^ level after refill was comparable to that before BHQ-induced depletion in STIM1-overexpressing cells (Figure 6G); however, the level of ER Ca^2+^ after refill was significantly higher than that before depletion in STIM1-TRNN-overexpressing cells. These results indicate that EB1 binding constitutes a mechanism that optimizes SOCE and prevents ER Ca^2+^ store overload.

## Discussion

Based on our findings, we propose a model in which EB1-mediated diffusion trapping of STIM1 optimizes SOCE at ER-PM junctions and prevents ER Ca^2+^ overload (Figure 7). In the resting state, EB1 binding traps STIM1 at growing MT ends, restricting the amount of freely diffusible STIM1 in the ER membrane. Following ER Ca^2+^ store depletion, EB1 binding continues to serve as a counterbalance mechanism, limiting STIM1 localization at ER-PM junctions and activation of SOCE. In subcellular regions without growing MT, freely diffusible STIM1 molecules in the ER can readily bind to PM PIP2 to mediate SOCE at ER-PM junctions following ER Ca^2+^ store depletion. Without the counterforce provided by EB1 binding, STIM1 stays longer at ER-PM junctions, resulting in elevated SOCE and ER Ca^2+^ store overload.

**Figure 7.**
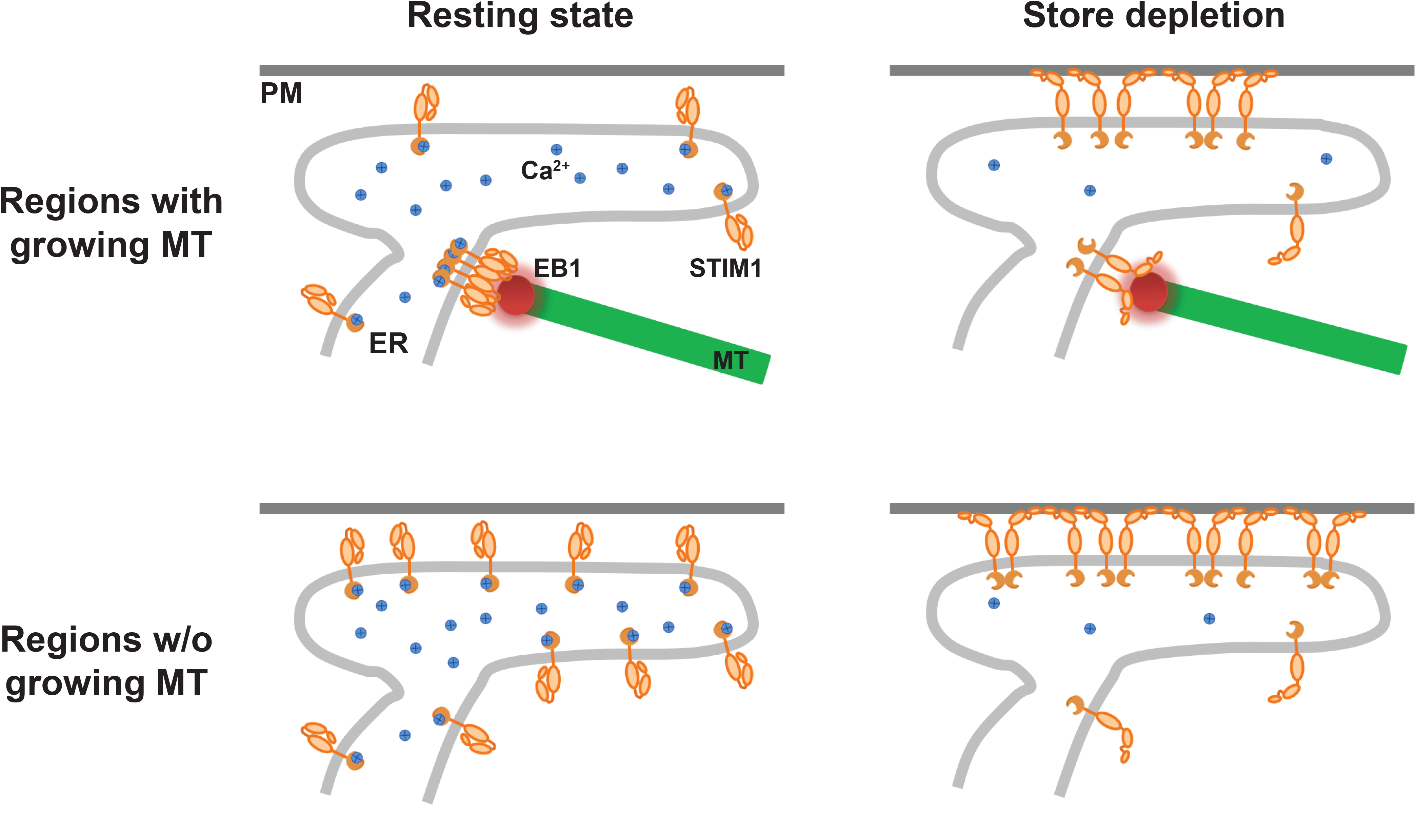
Model of STIM1-EB1 Interaction Regulates STIM1 Translocation to ER-PM Junctions.

This model can be further applied to understand physiological functions in cells with polarized MT distribution. During directed cell migration, MT plus-ends are orientated towards the front end of cells (Rodriguez et al., 2003). STIM1-EB1 interaction leads to polarized STIM1 distribution accompanied by low cytosolic and ER Ca^2+^ levels, indicative of limited SOCE, at the front end of migrating cells (Tsai et al., 2014). Additionally, disrupting the polarized distribution by introducing STIM1-TRNN abolished cell migration, demonstrating the importance of STIM1-EB1 interaction in STIM1 distribution and in maintaining polarized Ca^2+^ signaling for cell migration.

STIM1 translocation to ER-PM junctions is a complex process involving in a series of signaling events with multiple mechanisms contributing to STIM1 targeting to ER-PM junctions, such as STIM1-PIP2 binding and STIM1-Orai1 interaction (Prakriya and Lewis, 2015). Moreover, several STIM1/Orai1-interacting proteins, including SARAF (Palty et al., 2012), septins (Sharma et al., 2013), junctate (Srikanth et al., 2012), and CRACR2A (Srikanth et al., 2010), have been shown to regulate STIM1 translocation, STIM1-Orai1 interaction, or SOCE. These factors likely obscured previous studies in understanding of how STIM1-EB1 interaction contributes to SOCE and caused contradictory results (Baba et al., 2006; Bakowski et al., 2001; Ribeiro et al., 1997; Smyth et al., 2007). By generating iMAPPER-633, a mini-STIM1 construct that contains the ER targeting motifs, an FP, a chemically-inducible oligomerization unit, cytosolic linkers, and the CT of STIM1, the subcellular localization of STIM1 in the resting state and during ER Ca^2+^ depletion was successfully recapitulated. This reconstitution approach revealed that the EB1 trapping mechanism is dominant over PM targeting via the PB even in the exposed CT. We further found that the PB in the exposed CT is sufficient for PM targeting since disrupting STIM1-EB1 interaction led to a clear shift of iMAPPER-633 localization to ER-PM junctions without oligomerization. Oligomerization likely creates a stronger polybasic motif for PIP2 binding, enabling a shift of iMAPPER-633 trapping at MT plus-ends to ER-PM junctions. Thus, it is likely that STIM1 activation not only exposes the PB but also generates a stronger PB by oligomerization that counterbalances the EB1-mediated trapping mechanism, leading to translocation to ER-PM junctions. Consistent with this notion, STIM1-2K with an enhanced PB shifts the balance toward PM binding and distributes to ER-PM junctions prior to STIM1 activation.

We demonstrated that EB1 binding significantly reduces STIM1 diffusion in the bulk of the ER. This observation may explain the slow diffusion coefficient of STIM1 (~0.1 μm^2^/s) in the resting state (Liou et al., 2007; Wu et al., 2014); whereas ER membrane proteins in general have a diffusion coefficient of 0.2-0.5 μm^2^/s (Lippincott-Schwartz et al., 2000). Interestingly, single molecule tracking of STIM1 revealed a broad range of diffusion coefficients (Wu et al., 2014), which may represent a mixed population of EB1-free and EB1-trapped STIM1. Consistent with a previous study, we found that ionomycin-induced STIM1 translocation to ER-PM junctions is a much slower process with a t1/2 ~50 s compared to STIM1 oligomerization that occurs almost instantly during ER Ca^2+^ depletion (Liou et al., 2007). STIM1-TRNN showed a significantly faster translocation with ~2-fold increase in t1/2. The enhanced STIM1-TRNN translocation to ER-PM junctions further led to an accelerated Orail accumulation at ER-PM junctions, indicating that EB1 binding regulates the kinetics of SOCE.

Moreover, we demonstrated that the trapping mechanism mediated by EB1 works continuously during SOCE since activated STIM1 was recaptured by EB1 following the disruption of PM trapping by ML-9, a potent inhibitor of myosin light chain kinase (MLCK). The effects of ML-9 on STIM1 appeared to be independent of MLCK inhibition since knockdown of MLCK had no effect on SOCE (Smyth et al., 2008). Consistent with a previous observation that ML-9 was ineffective in inhibiting SOCE when both STIM1 and Orai1 were overexpressed, we found STIM1-Orai1 clusters remained similar following ML-9 treatment (Figure S2B). Thus, it is possible that ML-9 affects STIM1 interaction with PIP2 at the PM. There are multiple proteins localize at ER-PM junctions by binding to PM lipids to provide interorganelle signaling (Chang and Liou, 2016; Henne et al., 2015). Further work in defining the mechanisms underlying the actions of ML-9 may shed new light on STIM1 targeting and the function and regulation of ER-PM junctions.

A previous study demonstrated that STIM1 phosphorylation at residue S575, S608, and S621 by ERK1/2 is important for STIM1 dissociation from EB1 and translocation to ER-PM junctions during ER Ca^2+^ store depletion (Pozo-Guisado et al., 2013). Intriguingly, STIM1 phosphorylation was detected 2-5 min after TG treatment, arguing that STIM1 phosphorylation may occur after its translocation to ER-PM junctions. Nonetheless, STIM1 phosphorylation may provide a mechanism to disengage EB1 trapping for enhancing SOCE under certain conditions, such as cell migration (Casas-Rua et al., 2015). Phosphorylation of STIM1 may also be relevant during cell division since dissociation of phosphorylated STIM1 from EB1 is required for exclusion of the ER from mitotic spindles (Smyth et al., 2012).

SOCE is one of the most important pathways for Ca^2+^ signaling and homeostasis. Thus, the precise spatial-temporal regulation of SOCE is crucial for supporting cellular functions and health. Here we reveal an unexpected role of MT plus-ends in optimizing STIM1 translocation and SOCE via the EB1-mediated trapping mechanism and show that STIM1-EB1 interaction is important for preventing Ca^2+^ overload, which has been associated with many pathological conditions including stroke, neurodegeneration and cancer (Dong et al., 2006; Orrenius et al., 2003; Trump and Berezesky, 1995). Our study on the crosstalk between MT plus-ends and STIM1-mediated SOCE may shed light on how cells dynamically coordinate MT growth to regulate Ca^2+^ signaling in physiological processes.

## Materials and methods

### Reagents

Thapsigargin (TG), Pluronic F-127, and Fura-2 AM were purchased from Invitrogen (Carlsbad, CA). All chemicals for extracellular buffer (ECB, 125 mM NaCl, 5 mM KCl, 1.5 mM MgCl2, 20 mM HEPES, 10 mM glucose, and 1.5 mM CaCl2, pH 7.4), penicillin and streptomycin solution, ML-9, nocodazole (NocZ), ionomycin, histamine, and EGTA were obtained from Sigma (St. Louis, MO). AP20187 was purchased from Clontech (Mountain View, CA). BHQ was obtained from Calbiochem (Billerica, MA). Anti-EB1 antibody (ab53358) and anti-beta actin antibody (ab8227) were obtained from Abcam (Cambridge, MA). Human cDNA library and small interfering RNA (siRNA) used in this study were generated as described previously (Liou et al., 2005). Primers used for siRNA generation are listed in Table S1.

### Cell Culture and Transfection

HeLa cells purchased from ATCC (Manassas, VA) were cultured in MEM supplemented with 10% FBS (HyClone, Logan, UT) and penicillin and streptomycin solution. DNA plasmids (25-50 ng) and siRNAs (25 nM) were transfected into HeLa cells with TransIT-LT1 reagent for 16-20 hours and TransIT-TKO reagent for 48-72 hours, respectively (Muris Bio, Madison, WI).

### DNA Constructs

YFP-STIM1, YFP-STIM1-D76A, MAPPER, mCherry-STIM1, mCherry-ER (KDEL), Orai1-mCherry, mCherry-TM and mCherry-K Ras tail were described previously (Chang et al., 2013; Liou et al., 2007; Liou et al., 2005). mCherry-STIM1-D76A was constructed by replacing the YFP portion of YFP-STIM1-D76A with mCherry. iMAPPER-633 was generated by inserting PCR fragments of (i) 2X FKBP, (ii) TM plus cytosolic regions of MAPPER (without PM targeting motif), and (iii) STIM1 CT containing amino acid 633 to 685, into the MAPPER (YFP or mCherry) construct digested with SpeI and BamHI. CT mutants of STIM1 and iMAPPER-633 were generated using QuickChange site-direct mutagenesis kit (Agilent Technologies, Santa Clara, CA). STIM1-2K was generated by site-directed mutagenesis to insert a fragment encoding glycine, alanine, glycine and amino acids 671 to 685 before the stop codon of STIM1. EB1-GFP and EB1-mCherry were cloned by inserting a PCR fragment containing EB1 into GFP-N1 and mCherry-N1 plasmids, respectively. All constructs listed here were verified by sequencing. All oligonucleotides used in this study are listed in Table S1.

### Immunoprecipitation

HeLa cells were cultured on 6-well plates and transfected with EB1-GFP (300 ng/well) and mCherry-STIM1 subtypes (200 ng/well) for overnight. Cells were then washed with warm PBS before lysis with 20 mM Tris buffer, pH 7.5 containing 100 mM NaCl, 0.5% NP-40, and protease inhibitors on ice for 30 min. The lysates were subjected to centrifugation at 16,000 X g for 15 min at 4°C and the clear lysates (supernatants) were collected. The lysates were mixed with GFP-nAb agarose resin (Allele Biotechnology, San Diego, CA) and incubated with tumbling for 2 h at 4°C. The immunoprecipitated proteins were eluted with NuPAGE LDS sample buffer (Life Technologies, Carlsbad, CA) after washing the GFP-nAb agarose resin with 10 mM Tris buffer containing 150 mM NaCl and 0.5% NP-40 for four times. The eluted proteins were analyzed by Western blotting using antibodies against GFP (Abcam, ab290) or STIM1 (Cell Signaling Technology, #4916).

### Live-Cell Confocal and TIRF Microscopy and Image Analysis

HeLa cells were cultured and transfected on Lab-Tek chambered #1 coverglass (NUNC, Rochester, NY). Before imaging, cells were washed with ECB. Live-cell confocal and TIRF imaging experiments were performed at room temperature with a CFI Apo 60 X or 100 X objectives (NA 1.49) and a confocal-TIRF microscope custom-built using a Nikon Eclipse Ti microscope (Melville, NY) with an HQ2 camera and an EM camera (c9100-13; Hamamatsu). The microscope was controlled by Micro-Manager software (Edelstein et al., 2010). Fluorescence recovery after photobleaching (FRAP) experiment was performed with a 60 X objective and Andor spinning disk confocal and FRAPPA units on a Nikon Eclipse Ti microscope controlled by MetaMorph software (Molecular Devices, Sunnyvale, CA). A 4.65 μm^2^ area of HeLa cells expressing YFP-STIM1 or YFP-STIM1-TRNN were subjected to photobleaching with 3 pulses of 515 nm laser for 300 μs at maximal intensity. The intensity traces of the photobleached areas were analyzed and normalized to a non-bleached area in the same cells. For the analyses of STIM1 subtypes and Orai1 translocation to ER-PM junctions, 20 to 30 puncta in each cell from TIRF images were selected. The intensity traces of the selected puncta from the same cell were background subtracted, normalized to time zero, and averaged.

### Cytosolic and ER Ca^2+^ Levels Measurement

For measuring cytosolic Ca^2+^ levels, HeLa cells were loaded with 0.5 μM fura-2 AM in ECB containing 0.05% pluronic F-127 and 0.1% of BSA for 30 minutes at room temperature avoiding light. Loaded cells were then washed with ECB containing 0.1% BSA, and incubated in ECB for another 15-30 minutes before the experiments. Single-cell Ca^2+^ images were taken with a Plan Fluor 4X objective (NA 0.15) and an automated microscope custom-built on a Nikon Eclipse Ti microscope with a camera (HQ2; Photometrics). The microscope was controlled by Micro-Manager software (Edelstein et al., 2010). Intracellular Ca^2+^ levels were indicated by ratio of emission 510 nm excited at 340 nm over those at 380 nm. To measure ER Ca^2+^ levels, HeLa cells were co-transfected with D1ER and mCherry-STIM1 or mCherry-STIM1-TRNN. Single-cell Ca^2+^ images were taken with a Plan Fluor 40X objective (NA 1.30) and a confocal-TIRF microscope custom-built using a Nikon Eclipse Ti microscope with an HQ2 camera and an EM camera (c9100-13; Hamamatsu). Dynamic changes in ER Ca^2+^ levels were indicated by the ratio of FRET (CFP excitation-YFP emission) signal to that of CFP.

### Statistical Analysis

Data were statistically analyzed by t-test or one-way analysis of variance (ANOVA) using SigmaPlot software (Systat Software Inc., San Jose, CA).

## Acknowledgements

We thank Carlo Quintanilla for carefully reading the manuscript and the Liou laboratory members for valuable discussions and technical assistance. We also thank the University of Texas Southwestern Live-Cell Imaging Facility for assistance with FRAP experiments. We are grateful to Linda Patterson for administrative assistance. This work was supported by National Institutes of Health grant GM113079, Welch Foundation Grant I-1789, Howard Hughes Medical Institute Graduate Grant 56006776 (to the Mechanisms of Disease and Translational Science Ph.D. Track), Taiwan National Science Council Grant 102-2917-I-564-019. J. Liou is a Sowell Family Scholar in Medical Research. The authors declare no competing financial interests.

## Author Contributions

C.-L.C. and J.L. designed iMAPPER-633. C.-L.C., Y.-J. C., and J.L. performed experiments. C.-L.C. and J.L. analyzed the results. J.L. conceived and supervised the project. C.-L.C. and J.L. wrote the manuscript.

## Abbreviations

CT,: C-terminal region;
EB1,: end binding protein 1;
ECB,: extracellular buffer;
EF-SAM,: EF hand-sterile α motif;
ER,: endoplasmic reticulum;
FKBP,: FK506-binding protein; iMAPPER, inducible MAPPER;
IP3,: inositol 1,4,5-triphosphate;
MAPPER,: membrane-attached peripheral ER;
MT,: microtubule;
NocZ,: nocodazole;
PB,: polybasic motif;
PM,: plasma membrane;
PIP2,: phosphatidylinositol 4,5-bisphosphate;
SERCA,: sarco/endoplasmic reticulum Ca^2+^-ATPase
SOCE,: store-operated Ca^2+^ entry;
STIM1,: stromal interaction molecule 1;
TG,: thapsigargin;
TIRF,: total internal reflection fluorescence

## Supplemental Figure Legends

**Figure S1.**
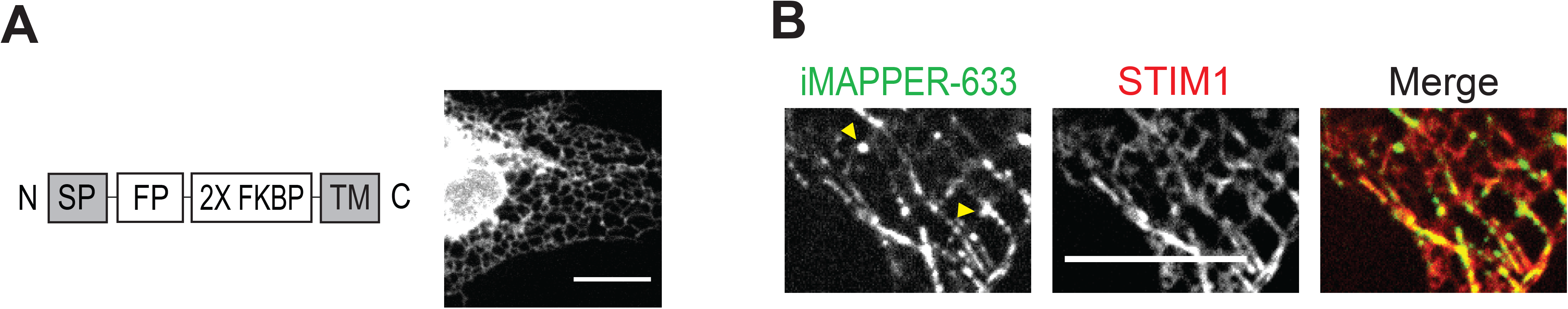
Localization of 2X FKBP-TM and iMAPPER-633. (A) A diagram of 2X FKBP-TM construct (left) and its ER localization as shown by confocal microscopy in HeLa cells. Scale bar, 10 μm. (B) The localization of YFP-iMAPPER-633 in HeLa cells coexpressing mCherry-STIM1 monitored by confocal microscopy. Yellow arrowheads indicate iMAPPER-633 puncta without STIM1 colocalization, possibly formed due to loss of EB1 binding after MT catastrophe. Scale bar, 10 μm.

**Figure S2.**
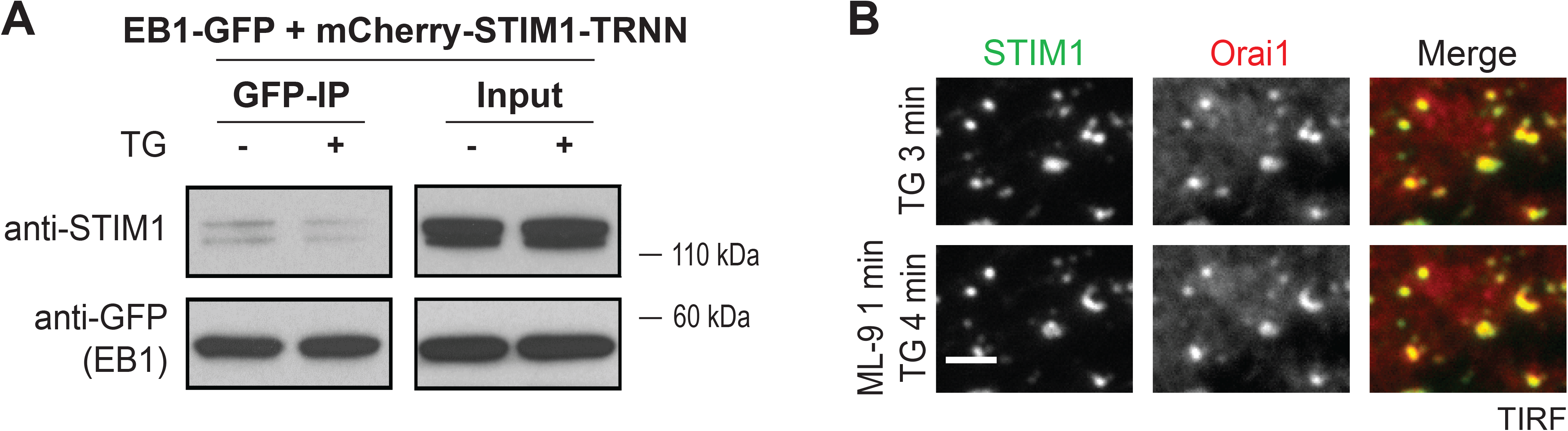
Minimal Interaction between STIM1-TRNN with EB1 and STIM1-Orai1 Complexes Localized at ER-PM Junctions following ML-9 Treatment. (A) Immunoprecipitation (IP) of EB1-GFP with mCherry-STIM1-TRNN following 1 μM TG treatment in HeLa cells. Protein levels of EB1-GFP and mCherry-STIM1-TRNN in total cell lysates (Input) and in IP were assessed by western blotting using antibodies against GFP (anti-GFP) and STIM1 (anti-STIM1). (B) YFP-STIM1 and Orail-mCherry puncta remain unchanged following 100 μM ML-9 treatment monitored by TIRF microscopy. Scale bar, 2 μm.

**Figure S3.**
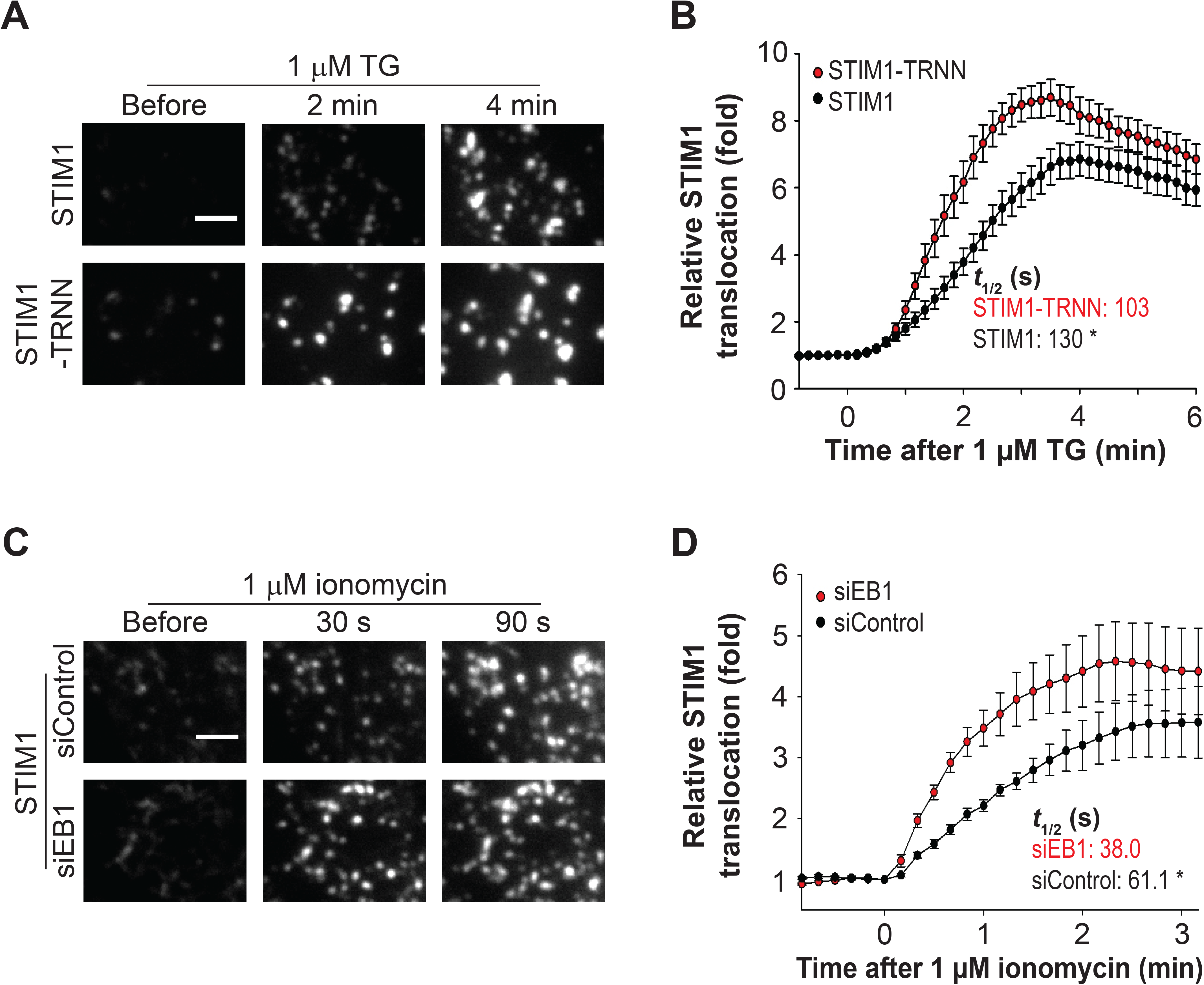
EB1 Binding Impedes STIM1 Translocation to ER-PM Junctions during ER Ca^2+^ Depletion. (A) Translocation of YFP-STIM1 and YFP-STIM1-TRNN to ER-PM junctions following 1 μM TG treatment in HeLa cells monitored by TIRF microscopy. Scale bar, 2 μm. (B) Relative translocation to ER-PM junctions of YFP-STIM1 and YFP-STIM1-TRNN as described in (A). Mean ± SEM are shown (19 to 22 cells from 4 independent experiments). Mean times to the half-maximal translocation (*t_1/2_*) are indicated. *, *p* < 0.05. (C) Translocation of YFP-STIM1 to ER-PM junctions following 1 μM ionomycin treatment in HeLa cells transfected with siControl or siEB1 monitored by TIRF microscopy. Scale bar, 2 μm. (D) Relative translocation to ER-PM junctions of YFP-STIM1 as described in (C). Mean ± SEM are shown (9 to 10 cells from 3 independent experiments). *t_1/2_* are indicated. *, *p* < 0.05.

Supplemental **Movie 1. iMAPPER-633 Binds to EB1 at MT Plus-ends.** iMAPPER-633 displayed comet-like structures moving toward the cell periphery monitored by confocal microscopy in HeLa cells transfected with YFP-iMAPPER-633. Scale bar, 10 μm.

Supplemental **Movie 2. Activated STIM1 Retains EB1 Binding Ability.** Activated form YFP-STIM1-D76A (green) trapped by EB1-mCherry (red) following 100 μM ML-9 treatment in HeLa cells monitored by confocal microscopy. Scale bar, 10 μm.

**Table S1.**
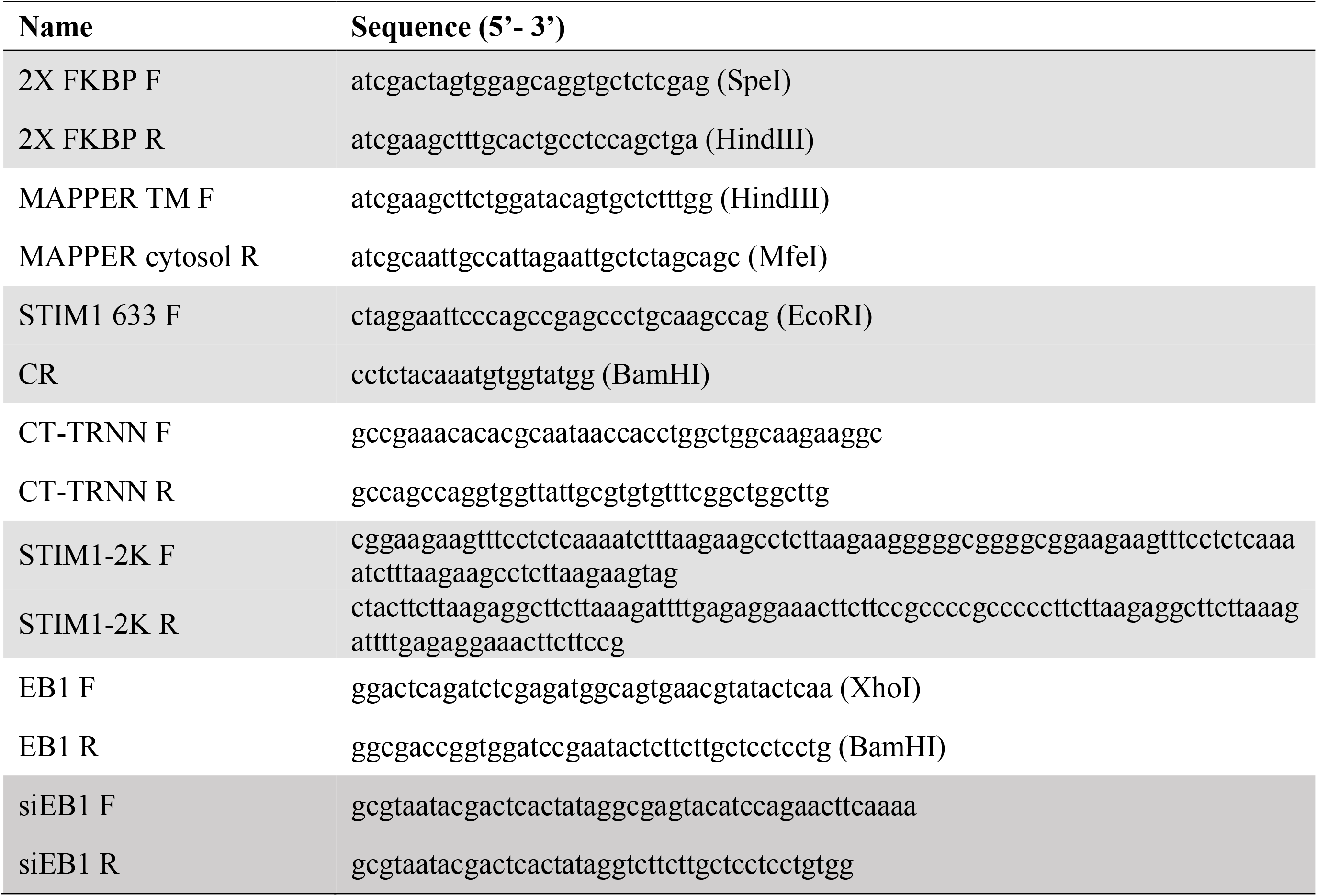
Oligonucleotides used in this study.

